# Novel Library Assembly Technique for Developing Nanobodies Targeting IPNv VP2 Protein

**DOI:** 10.1101/2024.11.13.623509

**Authors:** Camila Pino-Belmar, Johanna Himelreichs, Camila Deride, Tamara Matute, Isaac Nuñez, Severine Cazaux, Fernan Federici, Karen Moreno, Genaro Soto Rauch, Jose Munizaga, Denise Haussmann, Alejandro Rojas-Fernandez, Figueroa V Jaime, Guillermo Valenzuela-Nieto

## Abstract

Infectious pancreatic necrosis virus (IPNv) poses a significant threat to the global salmon farming industry, causing high mortality and economic losses. Given the limitations of traditional antibody therapies in aquaculture, we explored the development of nanobodies targeting the VP2 protein of IPNv. We developed a novel nanobody library generation method using a streamlined assembly protocol based on Type IIS restriction enzymes. This method enabled quick and efficient bacterial display library generation, and concatenated, multi-level cloning through an ingenious design in which the ligation of the first level generates a new restriction site that can be utilized in the subsequent cloning level, allowing for rapid transfer to other vectors. By combining this assembly approach with density gradient-based enrichment and high-throughput screening, we identified nanobodies that specifically recognize IPNv’s VP2 protein. Notably, the P9 clone demonstrated high specificity for IPNv in immunofluorescence assays, highlighting its diagnostic potential. Our method not only accelerates nanobody library generation but also enhances its quality and adaptability, marking a significant advancement in enhancing the responsiveness of nanobody development platforms against emerging pathogen outbreaks.

## Introduction

Infectious pancreatic necrosis virus (IPNv) poses a critical and persistent challenge for the global salmon farming industry. It is a double-stranded RNA virus belonging to the family Birnaviridae, genus Aquabirnavirus (Dobos, 1995). The genome of IPNv consists of two segments of double-stranded RNA, packaged within a non-enveloped icosahedral capsid approximately 60 nm in diameter. Segment A encodes the VP2, VP3, VP4, and VP5 proteins, while segment B encodes an RNA polymerase (VP1). VP2 and VP3 are the two main structural proteins of this virus (Azad et al., 1987; Dobos, 1995; Håvarstein et al., 1990). VP2 is a 54 kDa protein that is the primary component of the capsid by number of units and is responsible for binding to the host cell (Coulibaly et al., 2005, 2010; Dobos, 1995). It is composed of three domains: a central domain (S), a base located towards the interior of the viral particle, and an outward-facing spike or projection domain (P). The spikes, organized as VP2 trimers around an axis, contain the primary viral antigenic sites, the cell specificity epitope, and some virulence determinants. Since the 1980s, neutralizing epitopes in VP2 have been described, primarily located in an internal region of VP2 between amino acids 206 and 350, which is recognized by the monoclonal antibody (MAb) 17/82. This antibody has been reported to neutralize viral infection (Azad et al., 1987; Caswell-Reno et al., 1986; Håvarstein et al., 1990; Heppell et al., 1995). The host cell binding site is in the P domain of VP2, at the top of the spike; however, certain amino acids located in the groove of the spike, near the S domain, are also important for cell binding. It has been suggested that the former are responsible for recognition and binding, while the latter are involved in the internalization process (Coulibaly et al., 2010).

IPN virus primarily affects salmon during their freshwater stage, leading to extremely high mortality rates, reaching up to 90% in severe cases, especially among fry and juveniles (Asche and Bjørndal, 2011). High mortality not only disrupts the later development of fish in seawater but also has a significant economic impact. Surviving fish serve as viral reservoirs, thereby facilitating its spread in high-density aquaculture environments, such as recirculating aquaculture systems (RAS) and open-sea cages(Smail and Munro, 2012). Additionally, the virus can remain active in the environment, including in soil and aquaculture infrastructure, for several weeks, further complicating its control (Labraña et al., 2008). The economic losses associated with IPNv extend beyond direct mortality; the virus also induces epigenetic modifications that predispose fish to subsequent viral and bacterial infections during their seawater growth phase, significantly increasing losses by affecting both the quality and quantity of the final product (Manríquez et al., 2023).

Despite efforts to control IPNv through vaccines and selective breeding programs, there has been an observed increase in outbreak frequency, even in salmon populations genetically selected for IPNv resistance. Recent outbreaks among genetically resistant salmon in Chile, Scotland, and Norway suggest that the virus is developing mutations, particularly in the VP2 protein, enabling it to evade host defenses (Godoy et al., 2022; Hillestad et al., 2021). These outbreaks may be due to mutations in the hypervariable region (HPR) of VP2, which warrants further investigation (Godoy et al., 2022).

Emerging IPNv strains, characterized by mutations in the VP2 protein, have shown adaptability to different salmonid species and the ability to overcome immune barriers. This highlights the need for new therapeutic strategies, with immunotherapy presenting a promising advanced tool against viral variants. However, the high costs of monoclonal antibodies limit their feasibility for aquaculture. This is where a promising alternative emerges: the use of nanobodies. Derived from single-chain antibodies in camelids, nanobodies offer significant advantages over conventional therapies. Their small size and unique structure allow them to bind to viral epitopes that are inaccessible to other antibodies, resulting in high specificity and neutralizing capacity. Additionally, nanobodies are more stable and easier to produce than traditional antibodies, making them a cost-effective tool for scenarios that require affordable solutions. The use of nanobodies could be a flexible and effective solution to tackle the constant emergence of new variants, becoming a promising therapeutic tool to improve salmon health and reduce related economic losses (Jovčevska and Muyldermans, 2020; Wang et al., 2021).

Traditionally, a key step in identifying new nanobodies is generating a library from which subsequent screening can be performed to identify the best candidates. This is typically done using traditional cloning with restriction enzymes, following protocols that are often time-intensive to ensure optimal library quality. In this study, we explored alternative assembly methods to improve the efficiency of this process.,Due to their capacity to generate sequence variants in a single reaction, assembly techniques based on Type IIS enzymes, like the Golden Gate method and its derivatives, have proven highly effective for large-scale DNA library construction (Casini et al., 2015; Engler et al., 2014; Sarrion-Perdigones et al., 2013). These techniques enable the simultaneous assembly of multiple DNA fragments by designing unique overhangs, facilitating the seamless and efficient creation of highly diverse sequence libraries (Ellis et al., 2011; Engler et al., 2008). This approach offers a significant advantage over traditional cloning methods, which often require multiple ligation and transformation steps for each DNA fragment, a process that is labor-intensive and error-prone (Sarrion-Perdigones et al., 2013).

In contrast, Type IIS-based assembly methods, such as Universal Loop (uLoop), reduce errors and accelerate the cloning process by allowing multiple fragments to be assembled in a single-step reaction (Andreou and Nakayama, 2018; Pollak et al., 2020). Furthermore, these techniques can be designed to create new restriction sites post-ligation, facilitating additional rounds of cloning in different vectors for expanded applications. Type IIS enzyme-based methods enable high-throughput DNA library generation, making them invaluable tools for applications like protein engineering, pathway optimization, and synthetic biology, where diverse DNA libraries are essential for exploring genetic variability efficiently (Andreou and Nakayama, 2018; Engler et al., 2008, 2014; Halleran et al., 2018; Iverson et al., 2016; Lin and O’Callaghan, 2018; Sarrion-Perdigones et al., 2013)(Xie et al., 2014; Engler & Marillonnet, 2014). This study presents an innovative nanobody library assembly technique based on the use of Type IIS restriction enzymes, applied to identify nanobodies targeting the VP2 protein of IPNv.

## Material and methods

### Expression and purification of VP2 protein

The nucleotide sequence of the VP2 protein was obtained from GenBank (accession number AY379740.1), synthesized, and cloned into the pHLTV vector using Gibson assembly. Specific primers were designed to amplify the VP2 coding sequence (fwd: CTTCCAGGGTGGATCCATGAACACAAACAAGGCAACCGC, rev: AGCCGGATCAAGCTTCGGTCTTTGTAGCGCCCTCCTGCGGCC) and the vector backbone (fwd: GAGGGCGCTACAAAGACCGAAGCTTGATCCGGCTGCTAACAAAG, rev: TTGTTTGTGTTCATGGATCCACCCTGGAAGTACAGGTTTTC). To reduce the probability of aggregation via electrostatic repulsion between highly charged soluble polypeptide,the construct was assembled to express VP2 fused with a 6xHis tag and a Lipoyl domain (from dihydrolipoamide acetyltransferase). Thus, solubility was enhanced and adequate time for correct folding was allowed.dditionally, these solubility enhancers might act as intramolecular chaperones by participating in native folding of the target proteins (Lebendiker and Danieli, 2014; Zou et al., 2008). The resulting construct was then transformed into E. coli strain BL21. A pre-inoculum was prepared by adding 10 μL of glycerol stock from a previously transformed E. coli BL21 strain to 15 mL of liquid LB medium supplemented with ampicillin. This culture was incubated overnight at 37°C with shaking. The next day, 100 mL of LB medium was inoculated with the overnight culture and incubated at 30°C for 2.5 hours with shaking. This culture was then used to inoculate 500 mL of LB medium, which was incubated at 37°C with shaking for an additional 2.5 hours. The bacteria were harvested by centrifugation at 3000 xg for 15 minutes at 4°C. The bacterial pellet was resuspended in 1 L of 2YT medium, incubated at 42°C for 30 minutes with agitation, and then cooled to 16°C for 10 minutes. The optical density at 600 nm (OD600) was measured to ensure a range between 0.6 and 0.8. Protein expression was induced by adding 10 μM IPTG, and the culture was incubated at 16°C for 40 hours with shaking. After induction, the bacteria were collected by centrifugation at 4000 rpm for 15 minutes at 4°C. Induced bacterial pellets from 1 L culture were resuspended in 10 mL of lysis buffer (50 mM Tris (pH 7.5), 500 mM NaCl, 10 mM imidazole, 1 mM DTT, 1X protease inhibitor). Cells were lysed by sonication (50% pulse, 20-second on/off intervals for 5 minutes) and centrifuged at 21,000 g for 45 minutes at 4°C. The supernatant was recovered and filtered through a 0.22 μm syringe filter. Protein purification was conducted using Ni-NTA affinity chromatography. The filtered lysate was applied to a Ni-NTA column containing 3 mL of pre-equilibrated agarose beads, compacted to a 2 mL bead bed. The column was washed with 20 mL of binding buffer (50 mM Tris (pH 7.5), 500 mM NaCl, 10 mM imidazole, 1 mM DTT), followed by 30 mL of wash buffer (50 mM Tris (pH 7.5), 500 mM NaCl, 30 mM imidazole, 1 mM DTT), and the target protein was eluted with 10 mL of elution buffer (50mMTris (pH 7,5), 500mM NaCl, 150mM imidazol, 1mM DTT). Purified protein was verified by SDS-PAGE followed by Coomassie staining.

### Alpaca Immunization

The alpaca immunization procedure followed the ethical guidelines of the Bioethics Committee of the Austral University of Chile (certifications 463/2022) and carried out as previously described by (Valenzuela Nieto et al., 2021). A 5 mL blood sample was collected one day before immunization to obtain pre-immune serum. For the initial immunization (day 1), 100 μg of recombinant VP2 protein was used. The recombinant protein elution buffer was exchanged to PBS 1X, then mixed with 2 mL of Veterinary Vaccine Adjuvant (GERBU FAMA) and administered subcutaneously to a male alpaca (*Vicugna pacos*) at four different sites, delivering a total volume of 4 mL. Seven days post-immunization, a 5 mL blood sample was taken.On day 14, a second immunization was performed using 100 μg of Spike protein. On day 15, a 120 mL blood sample was collected from the jugular vein into tubes containing 3.8% sodium citrate as an anticoagulant. This blood was diluted with an equal volume of HBSS medium (Gibco) without calcium, and the mixture was aliquoted into 10 mL portions, each layered on 5 mL of Ficoll-Paque Premium (GE Healthcare) in 15 mL sterile Falcon tubes. After centrifugation (1200 rpm, 80 min, room temperature), the PBMC layer was carefully collected, washed twice with PBS by centrifugation (3500 rpm, 10 min), and resuspended in 4 mL of sterile PBS (Gibco). RNA extraction was performed using the RNeasy Mini Kit (Qiagen), and cDNA synthesis was conducted with the QuantiTect Reverse Transcription Kit (Qiagen), following the manufacturer’s protocols.

### Library preparation

Both plasmids pNeae2 and pHEN6 (Salema and Fernández, 2013; Salema et al., 2016) for bacterial display libraries and periplasmic expression, respectively, were modified by mutagenic PCR in order to exchange the *SfiI* and *NotI* restriction sites for *SapI* and *BsaI* sites, based on the cloning strategy described by (Pollak et al., 2020) obtaining pNeae2.1 and pHen6.1.

Initial PCR amplification was performed using cDNA from immunized alpaca. The reaction mixture (50 μL) included 1 μL of 10 mM dNTPs, 10 μL of 5x Flexi GoTaq Buffer, 4.5 μL of 25 mM MgCl2, 2 μL of 10 μM primer CALL001 (GTCCTGGCTGCTCTTCTACAAGG), 2 μL of 10 μM primer CALL002 (GGTACGTGCTGTTGAACTGTTCC) (Conrath et al., 2001), 0.5 μL of GoTaq (5 U/μL), 2 μL of template DNA, and 28 μL of H2O. PCR conditions were as follows: initial denaturation at 94°C for 5 minutes, followed by 30 cycles of 94°C for 1 minute, 54°C for 1 minute, and 72°C for 1 minute, with a final extension at 72°C for 10 minutes. Products were run on a 1.5% agarose gel to isolate the 700 bp band, which was purified using the GenElute Gel Extraction Kit (Sigma), yielding approximately 100–200 ng/μL in 30 μL of preheated H2O.The 700 bp purified product was used as a template in a second PCR. The reaction mixture was similar to the first PCR, replacing primers CALL001 and CALL002 with GV145 (AAGCTCTTCATCCTATGGCTCAGGTGCAGCTGGTGG) and GV174 (TTGCTCTTCTTCGGAGGAGACGGTGACCTGGGTC), respectively. PCR conditions were 94°C for 5 minutes, followed by 30 cycles of 94°C for 1 minute and 72°C for 1 minute, with a final extension at 72°C for 10 minutes. The resulting 400 bp band was isolated, purified as described, and eluted in 30 μL of H2O.In a 0.2 mL PCR tube kept on ice, 1.5 μg each of vector pNeae2.1 and insert DNA were combined with 5 μL of 10x T4 Ligase Buffer (NEB), 5 μL of 10x rCutSmart Buffer, 5 μL of SapI (10 U/μL), 0.5 μL of T4 Ligase (2000 U/μL), and H2O to a final volume of 100 μL. The reaction was subjected to 40 cycles of 1 minute at 37°C, 1 minute at 16°C, followed by 10 minutes at 55°C, 10 minutes at 80°C, and held at 4°C.To the 100 μL reaction, 300 μL of cold 100% ethanol and 10 μL of 3M sodium acetate (pH 5.2) were added. After a 10-minute incubation at −20°C, samples were centrifuged at 20,000 x g for 30 minutes at 4°C, and the pellet was washed with 500 μL of cold 70% ethanol. The pellet was air-dried for 10 minutes and resuspended in 10 μL of preheated H2O at 70°C.SOC medium was pre-warmed to 37°C. Electroporation cuvettes (0.1 cm gap) were cooled at −80°C. Competent cells (50 μL aliquots) were mixed with 2.5–3.3 μL of DNA and electroporated at EC2 setting on a Bio-Rad Gene Pulser. Cells were recovered in 1 mL of SOC and incubated at 37°C with shaking at 200 rpm for 1 hour. Transformants were plated on LB agar with chloramphenicol and 2% glucose, incubated overnight at 37°C, and library size was estimated based on colony counts from serial dilutions. Protocol available at DOI: dx.doi.org/10.17504/protocols.io.q26g7mbwqgwz/v1

### Density gradient enrichment

This protocol was carried out according to (Valenzuela Nieto et al., 2021). NHS-activated Sepharose 4 Fast Flow beads (1 mL) were pre-washed with cold 1 mM HCl and sterile PBS, then incubated overnight with 200 μg of purified protein in PBS with rotation. Unreacted NHS groups were blocked with 0.5 M ethanolamine, and beads were washed with PBS and stored at 4°C. Library and control bacterial cultures were grown overnight in LB medium with chloramphenicol (25 μg/mL) and 2% glucose or kanamycin (50 μg/mL), respectively. Cultures were then diluted to OD600 = 0.02 in LB with antibiotic (without glucose) and grown to OD600 0.45–0.6 at 37°C, followed by induction with 50 μM IPTG at 30°C for 3 hours. Both library and control cultures were washed with PBS, adjusted to equal OD, and incubated with 300 μL of protein-coupled beads for 30 minutes at room temperature. The bead-bacteria mixture was layered onto 6 mL of Ficoll-Paque PLUS (GE Healthcare) in a 15 mL conical tube and centrifuged at 200 x g for 1 minute. The unbound fraction was discarded, and the bead pellet resuspended in 4 mL of PBS, rotated for 5 minutes at room temperature. This wash step was repeated six times. After washing, 1 mL of LB medium was added to the bead suspension and incubated for 5 minutes at room temperature. Aliquots of 50 μL were plated on LB agar with either 50 μg/mL kanamycin or 25 μg/mL chloramphenicol, both containing 2% glucose. The remaining sample was plated on additional LB chloramphenicol/glucose agar plates. Plates were incubated at 37°C overnight (20+ hours recommended). Colony counts from kanamycin and chloramphenicol first plates were used to assess the specific enrichment of Nanobody-expressing bacteria from the library.

### Candidate screening ELISA

Pools of 10 colonies or individual colonies obtained from density gradient separation were inoculated in 2 mL of LB medium and incubated overnight at 37°C with agitation at 200 rpm. A 100 μL aliquot of the overnight culture was transferred to 1.9 mL of fresh LB medium with 25 μg/mL chloramphenicol and incubated at 37°C with 200 rpm shaking until reaching an OD600 of 0.45–0.6. Protein expression was induced by adding IPTG to a final concentration of 50 μM for 3 hours at 30°C with 200 rpm shaking. Cultures were pelleted, resuspended in 1 mL PBS containing 0.2% Triton X-100, sonicated on ice for 10 seconds at 40% amplitude, and centrifuged at 14,000 x g for 30 minutes at 4°C. The supernatant containing the total protein extract was collected for each clone.

Ninety-six-well ELISA plates (Nunc MaxiSorp, ThermoFisher) were coated with 100 ng of full-length Spike protein or the receptor-binding domain (RBD) diluted in 1x PBS (pH 7.4) and incubated for 1 hour at 37°C. Plates were washed three times with PBS-T (PBS containing 0.05% Tween-20) for 5 minutes each and blocked with KPL blocking solution (Seracare) for 30 minutes at room temperature with gentle shaking. After discarding the blocking solution, each well was incubated with total protein extract from each clone or pool of clones diluted 1:5 in TBS-T with 5% BSA and incubated for 1 hour at room temperature with shaking. Plates were washed three times with PBS-T for 5 minutes each.The wells were then incubated with Mouse Anti-Myc antibody (9B11, Cell Signaling Technology) diluted 1:3000 in PBS-T with 5% BSA for 1 hour at room temperature, followed by three 5-minute washes with PBS-T. Subsequently, Goat Anti-Mouse IgG conjugated to HRP (Invitrogen) at a 1:5000 dilution in PBS-T with 5% BSA was added for 1 hour at room temperature, followed by three additional 5-minute washes with PBS-T. Signal development was achieved by adding 100 μL of 1-Step Ultra TMB-ELISA substrate (ThermoFisher) to each well and incubating for 15 minutes at 37°C, then for 5 minutes at room temperature. Finally, 100 μL of STOP Solution (ThermoFisher) was added, and absorbance was measured at 450 nm using a microplate reader.

### Subcloning and Transformation

Nanobody coding sequences were subcloned into the pHEN6.1 vector via a second round of Type IIs enzyme assembly using BsaI (NEB). In a 0.2 mL PCR tube on ice, 1.5 μg each of pHEN6.1 vector and purified plasmid DNA from selected nanobody clones were combined with 5 μL of 10x T4 Ligase Buffer, 5 μL of 10x rCutSmart Buffer, 5 μL of BsaI (10 U/μL), 0.5 μL of T4 Ligase (2000 U/μL), and H2O to a final volume of 100 μL. The reaction was cycled 40 times at 37°C for 1 minute and 16°C for 1 minute, followed by 10 minutes at 55°C, 10 minutes at 80°C, and held at 4°C. Following assembly, 5 μL of the reaction mixture was transformed into E. coli WK6 cells via heat shock at 42°C. After recovery in 1 mL SOC medium at 37°C with shaking (200 rpm) for 1 hour, transformants were plated on LB agar with chloramphenicol and 2% glucose, and incubated overnight at 37°C.

### Periplasmic Expression and Nanobody Purification

The E. coli WK6 strain was used for periplasmic expression using a modified protocol from (Salema and Fernández, 2013). A single colony of pHEN6-W25-transformed cells was cultured in 20 mL of LB medium with 100 μg/mL ampicillin and 1% glucose at 37°C with agitation for 16 hours. This pre-culture was diluted into 1 L of Terrific Broth (TB) containing 100 μg/mL ampicillin, 2 mM MgCl2, and 0.1% glucose, and grown at 37°C to an OD600 of 0.6–0.9. Nanobody expression was induced by adding 1 mM IPTG, followed by incubation at 28°C for 20 hours. Bacteria were harvested by centrifugation at 8000 rpm for 8 minutes at 4°C.The bacterial pellet was resuspended in 12 mL TES buffer (0.2 M Tris, 0.5 mM EDTA, 0.5 M sucrose, pH 8.0) and incubated on ice for 1 hour. An additional 18 mL of TES buffer was added, and the suspension was incubated for another hour on ice. Cell debris was pelleted by centrifugation at 8000 rpm at 4°C. The supernatant was applied to a 5 mL HisPur Ni–NTA agarose resin pre-equilibrated with binding buffer (50 mM Tris, 500 mM NaCl, 10 mM imidazole, pH 7.5). Bound proteins were washed with binding buffer (50 mM Tris, 500 mM NaCl, 10 mM imidazole, pH 7.5) and eluted in 15 mL of elution buffer (50 mM Tris, 150 mM NaCl, 150 mM imidazole, 1 mM DTT, pH 7.5). Purified Nanobodies were analyzed by SDS-PAGE to confirm purity and integrity.

### Immunofluorescence

Imprints of *Salmo salar* head kidney of IPNv infected and healthy individuals, tested by qPCR, were fixed with methanol 30min and then washed three times with PBS. Cells were incubated for 45 minutes at 37°C with purified Nanobodies. Cells were then washed three times in PBS, followed by incubation with a mouse anti-myc antibody (1:3000, Cell Signaling) for 45 minutes at 37°C. After three PBS washes, an anti-mouse Alexa 647 secondary antibody was applied for 30 minutes at 37°C. Nuclei were stained with 0.1 mg/mL DAPI for 10 minutes at room temperature. After a final PBS wash, samples were mounted using ProLong™ Gold Antifade Mountant (Thermo). Images were acquired using a Celldiscoverer 7 high-content automatic microscope (Carl Zeiss GmbH, Jena, Germany).

### Structure prediction

The structural prediction of the P9 nanobody was performed using AlphaFold, a machine learning model for protein structure prediction. The amino acid sequence of P9, including annotated complementarity-determining regions (CDRs), was input into the AlphaFold Protein Structure Database. AlphaFold-generated models were analyzed to confirm the overall folding and structural integrity, focusing particularly on the spatial orientation of CDR loops, which are critical for antigen recognition (Jumper et al., 2021). The resulting structure was visualized and annotated to highlight CDRs, facilitating further insights into the binding potential of P9 against the IPNv VP2 protein.

## Results

### Recombinant VP2 expression and purification

A 1.5 kb fragment encoding VP2 (GenBank accession number AY379740.1) was successfully cloned into the pHLTV vector using Gibson assembly, enabling expression with an N-terminal 6xHis tag and a dihydrolipoamide acetyltransferase (E2) lipoyl domain to enhance solubility (Fig. 1a). The resulting construct was transformed into E. coli strain BL21, followed by induction and purification using Nickel affinity chromatography. This process yielded 10 mL of recombinant VP2 protein at a concentration of 0.5 mg/mL, which were subjected to SDS-PAGE (Fig. 2b) examine the target protein expression. A band near 70kDa was detected consistent with expected size of 6xHis-Lipoyl domain-Tev site-VP2 fusion protein (66.571 kDa).

**Figure 1.**
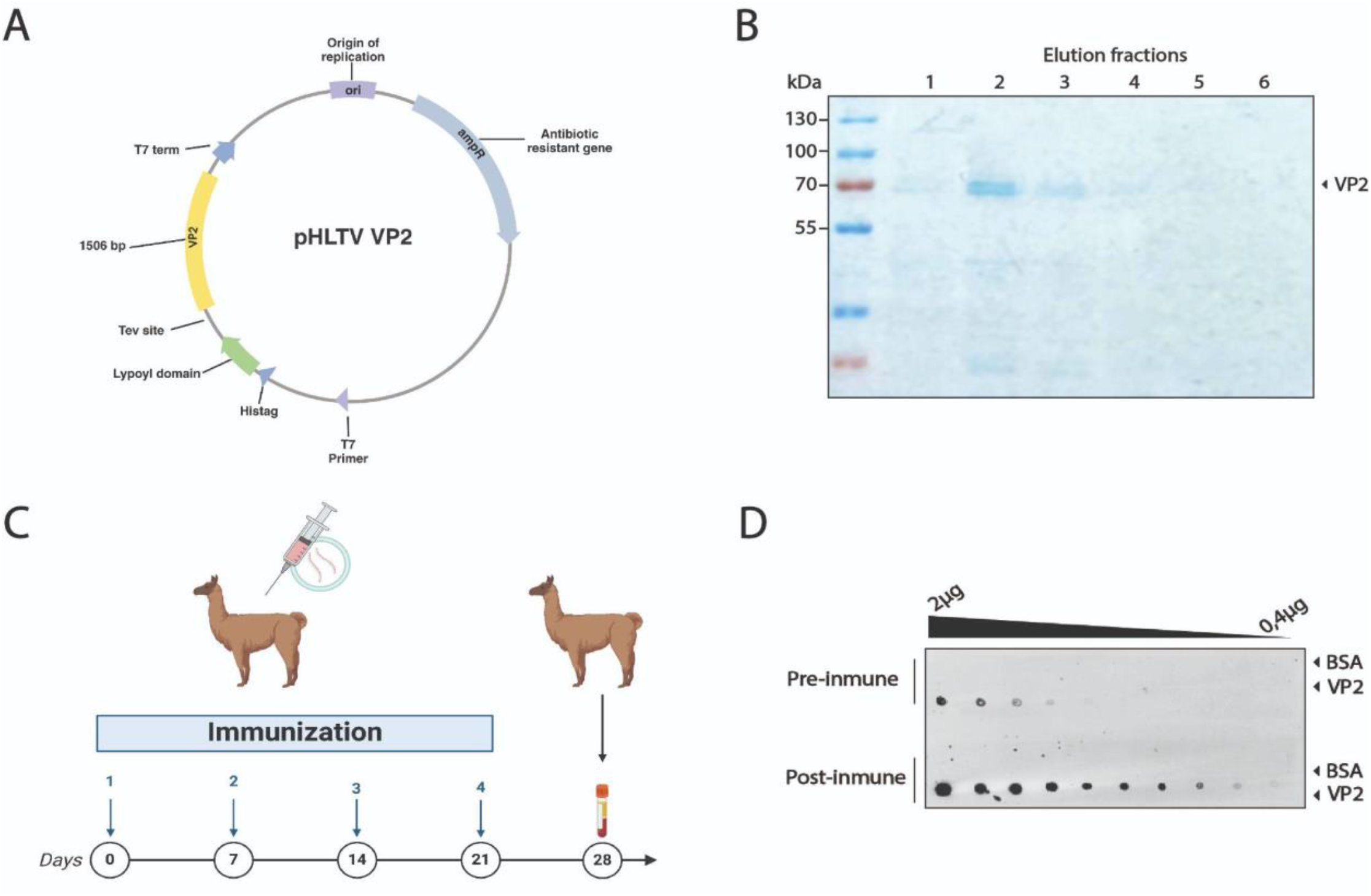
Recombinant VP2 protein production and immunization. (A) Scheme of the vector generated for the expression and purification of the IPNv VP2 protein in *E. coli*. The coding sequence for the protein, totaling 1506 bp, is highlighted in yellow. (B) SDS-Page to ensure protein integrity of recombinant VP2 protein before immunization. (C) Diagram of the alpaca immunization process. (D) Evaluation of the alpacas immune response by dot blot. Image shows the reaction to decreasing amounts of recombinant IPNv VP2 protein and bovine serum albumin (negative control) using a preimmunization control, and post immunization with VP2, using alpaca serums as a primary antibody source followed by an anti-camelid IgG-HRP secondary antibody.

**Figure 2.**
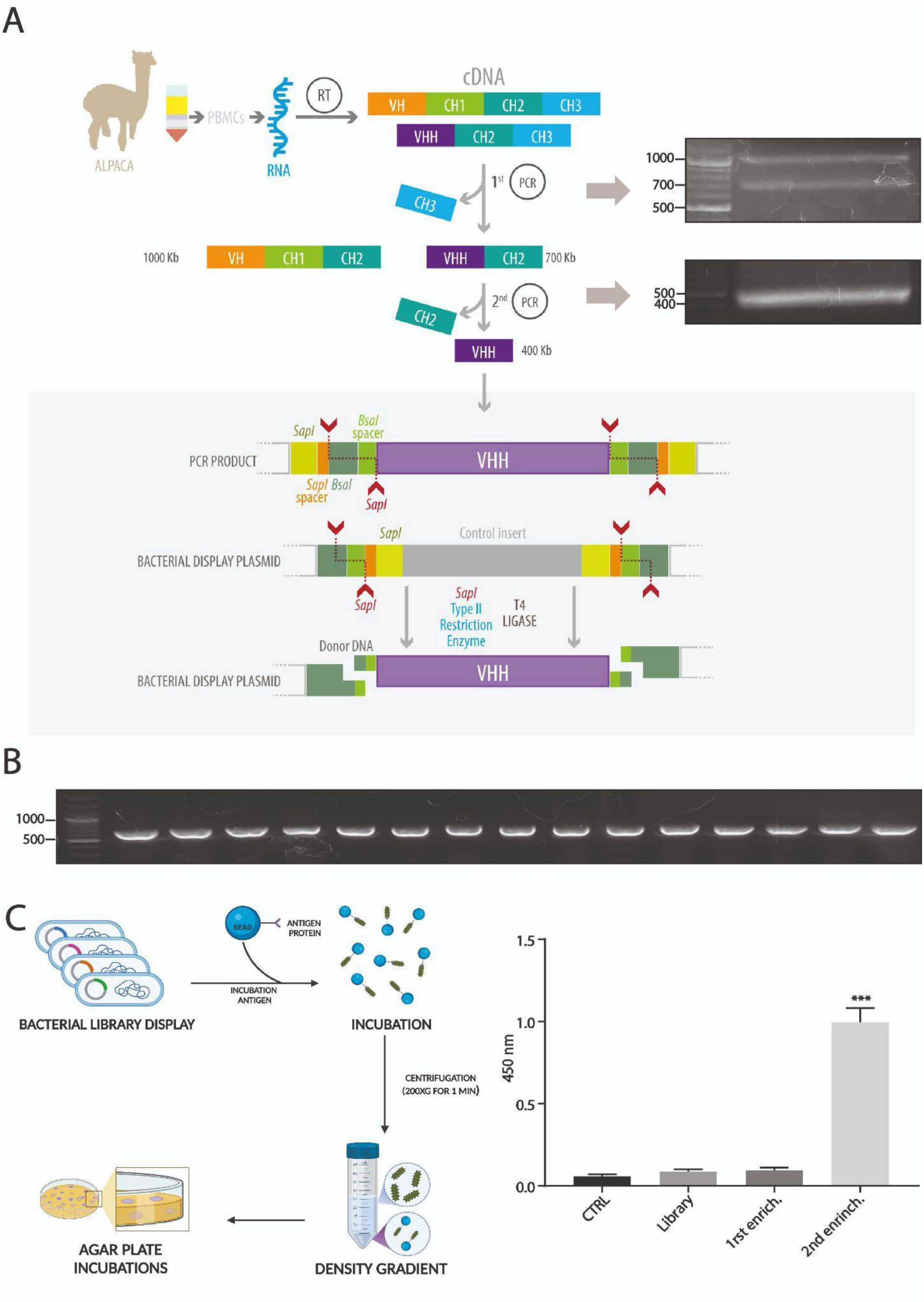
Novel Type IIS Restriction Enzyme-Based Method for Bacterial Display Library Generation. (A) Overview of the library construction process starting from RNA extraction from alpaca PBMCs, followed by reverse transcription to generate cDNA, and amplification of the VHH coding region through two rounds of PCR. The side panels display agarose gel electrophoresis of the PCR products obtained at each step during the library generation for VP2. The amplified VHH fragment is inserted into a bacterial display plasmid using *Sap*I and Type IIS restriction sites, allowing seamless assembly of the library through a one-tube digestion-ligation cyclic reaction that also generates a new restriction site, *Bsa*I. (B) Gel electrophoresis results confirming successful insertion and uniform size of VHH fragments in the bacterial display library clones. (A) and (B) electrophoresis are TAE 1X 1% agarose gels using Generuler 100bp plus standard (Thermo) (C) Schematic of the density gradient enrichment protocol, including incubation with beads coated in target antigens (VP2 recombinant protein) followed by a ficoll gradient-based separation. (D) ELISA assay of VP2 protein using total protein extract from library and the following density gradient enrichment rounds. This shows a significant signal increase after the second round, in which approximately 300 clones were recovered and used in subsequent screening. n = 3. Statistic t-test, *** P ≤ 0.001to control. Illustration (a) by Felipe G. Serrano BSc., MSc Scientific illustrator.

### Alpaca immunization

A male alpaca was immunized four times over a 28-day period with 100 μg of recombinant VP2 protein combined with an adjuvant (Fig. 1c). The animal health was monitored throughout the study via clinical examinations, hematological analysis, and serum biochemistry. The immune response in the alpaca’s serum was assessed using Dot blot analysis as a rapid qualitative method, immobilizing the antigen onto a nitrocellulose membrane and using the alpaca serum as the source of primary antibodies. An important increase of antigen specific IgG antibodies in the alpaca’s serum was observed, resulting in a post-immune serum capable of detecting even picograms very small amounts of the protein, such as those found in the tenth dilution, approximately 0,4ng (Fig-1d). Additionally, pre- and post-immunization sera were compared as primary antibodies to detect the viral protein in anterior kidney imprints of *Salmo salar* positive for IPNv infection. A clear increase in the signal was observed with the post-immune serum compared to the pre-immune serum and the same post-immune serum in IPNv-negative samples (Supplementary Fig. 1)

### Library preparation

The pNeae2 and pHEN6 plasmids were successfully modified by mutagenic PCR to replace the *Sfi*I and *Not*I restriction sites with *Sap*I and *Bsa*I, following the cloning strategy described by Pollak et al. (2020). The resulting plasmids, named pNeae2.1 and pHen6.1, enabled improved cloning compatibility for subsequent experiments. Our method for nanobody isolation builds on the bacterial display system previously described by Salema and collaborators (Salema et al., 2013), which has been successfully utilized by our team in earlier studies (Valenzuela Nieto et al., 2021). We implemented a novel procedure for library generation, utilizing Type IIs restriction enzymes that recognize specific DNA sequences but cut at distinct sites a defined number of base pairs away based on the uLoop system (Pollak et al., 2020). This approach involves an innovative strategy where cutting sites are arranged on both the insert and the vector in such manner that upon ligation, the *Sap*I recognition site is removed, enabling the reaction to proceed, all in one tube, in alternating cycles of digestion and ligation at 37°C and 16°C, respectively. As the *Sap*I sites on the vector are progressively eliminated with each ligation of the VHH insert (the coding region for the nanobody), the reaction increasingly favors the formation of vector-insert constructs, cutting dramatically the time needed to generate a bacterial display library (Fig. 2a). Ligation also assembles a new restriction site for *Bsa*I, useful for further cloning steps. The detailed protocol for library assembly is provided in the Materials and Methods section. Briefly, it involves two rounds of PCR: the first round amplifies the VH, CH1, and CH2 regions of heavy-chain antibodies, as well as the VHH and CH2 regions of single-chain antibodies, yielding bands of approximately 1 kb and 700 bp, respectively (Fig.2a first panel). These bands are separated electrophoretically, retaining only the 700 bp band for use as a template in a subsequent PCR reaction. This second PCR specifically amplifies the VHH coding region for nanobodies, adding *Sap*I sites and adaptors required for subsequent ligation into the pNeae2.1 vector, obtaining a PCR product around 400bp (Fig.2a second panel). Thus, we constructed a bacterial display library with a complexity of 1 × 10^6^ independent clones by electroporation of *E. coli* DH10B-T1R strain. Random clones were screened by PCR to verify insert size, with 100% of clones showing inserts of the expected size for a nanobody. This was further confirmed by sequencing, which additionally demonstrated that 100% of the clones corresponded to nanobodies and were in the correct reading frame, and with the *Bsa*I site in the right position (Fig. 2d).

We used a novel procedure for Nanobody selection using a simple Ficoll density gradient, previously used by our group with great success (Valenzuela Nieto et al., 2021). This takes advantage of the bacterial display system, in which each bacterium in the library expresses a single Nanobody clone (Salema et al., 2013). *E. coli* bacteria express intimin-Nanobody fusion proteins anchored in the outer membrane, exposing the functional Nanobody to the extracellular space for antigen recognition. Bacteria expressing specific nanobodies on their surface were incubated with Sepharose beads covalently coated with recombinant VP2, the antigen of interest, and migrated to the bottom of the Ficoll density gradient, while unbound bacteria remained in the upper fraction. Two rounds of enrichment were performed using this protocol, and the ability of the resulting clone pools from each round to recognize recombinant VP2 protein was assessed by ELISA. These enriched pools were also compared to the complete library and a control lacking nanobodies. A significant increase in VP2 detection by ELISA was observed for the clone pool obtained after the second round of density gradient enrichment, making this pool the optimal choice for identifying specific clones (Fig. 2e).

### Nanobody candidate selection

Following Nanobody selection using the density gradient protocol, approximately 300 colonies were obtained on LB-agar plates from the Sepharose-antigen coated fraction. We optimized conditions to extract intimin-Nanobody fusions from the bacterial outer membrane and used these protein extracts directly to assess binding to the VP2 protein by ELISA. Initially, 30 pools of 10 clones each were analyzed by incubating VP2-coated ELISA plates with bacterial extracts containing Nanobodies. Sequential incubation with a mouse anti-myc antibody, followed by an anti-mouse HRP-conjugate, revealed that 13 pools showed significant recognition of recombinant VP2. A secondary screening of individual clones was then performed (Fig. 3a). For example, in the pool containing clones 1-10, clone P9 demonstrated a strong ability to recognize recombinant VP2 in the ELISA assay (Fig. 3b). The selected clone was sequenced and CDR regions were identified using IMGT/V-QUEST (Brochet et al., 2008) (Fig. 3c).Finally, structure was predicted using AlphaFold, highlighting the CDRs (Fig. 3e and Suppl. Fig. 2).

**Figure 3.**
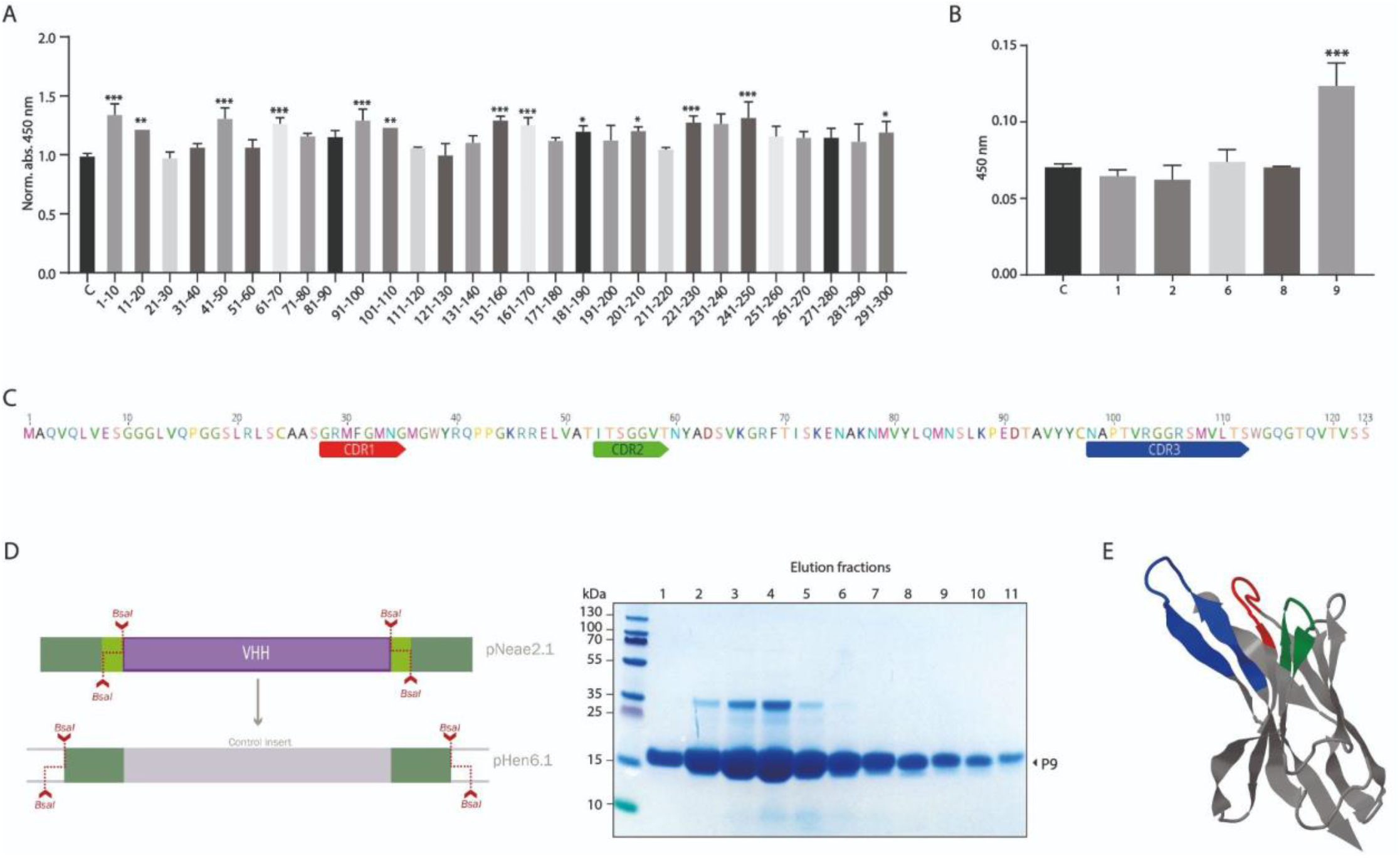
Identification and expression of a nanobody against IPNv VP2. (A) Initial screening of nanobody pools (30 pools, each containing 10 clones) was performed using ELISA on VP2-coated plates and direct total protein extracts of clones as primary antibodies. Mouse anti-Myc (1:3000) followed by anti-mouse-HRP were used for detection. Among these, 13 pools displayed significant binding to recombinant VP2 protein. Graphs depict the normalized to control mean of 450nm absorbance, n = 3. Statistic t-test, *** P ≤ 0.001; ** P ≤ 0.005; * P ≤ 0.01 to control (B) Example of secondary screening through ELISA of individual clones from 1-10 positive pool, in which we identified clone P9 as a positive binder for VP2. (C) Sequence analysis of clone P9, using IMGT/V-QUEST identified the complementary determining regions (CDRs), crucial for binding specificity. (D) Diagram of the subcloning strategy used to enable periplasmic expression of the isolated P9 clone. This strategy leverages the *Bsa*I site generated during library cloning, allowing a rapid transition to a new vector, in this case, pHEN6.1. Additionally, the SDS-PAGE on the right shows the purification of the P9 clone from the *E. coli* periplasm. (E) Alphafold structural prediction based on P9 sequence, CDRs are depicted in the same color scheme as in (C).

### Nanobody expression and validation

P9 clone was subsequently cloned into the pHen6.1 vector for periplasmic bacterial expression, using *Bsa*I sites and a similar assembly strategy than the one used for library construction (Fig. 3d). Large amounts of recombinant P9 Nanobody were obtained, around 20mg for liter of culture. (Fig. 3e). The purified P9 nanobody was used as primary antibody in immunofluorescence assays with imprints of *Salmo salar* head kidney samples, positive (qPCR tested) for IPNv. We observed a characteristic fluorescence pattern, consistent with previous reports using commercial antibodies (Abdullah et al., 2015; Levican et al., 2017), with no signal detected in negative or uninfected controls. This suggests that the P9 nanobody specifically recognizes IPNv-infected cells (Fig.4). Thus, our experiment suggests the P9 Nanobody is applicable as a tool for the direct diagnosis of infected cells by immunofluorescence.

**Figure 4.**
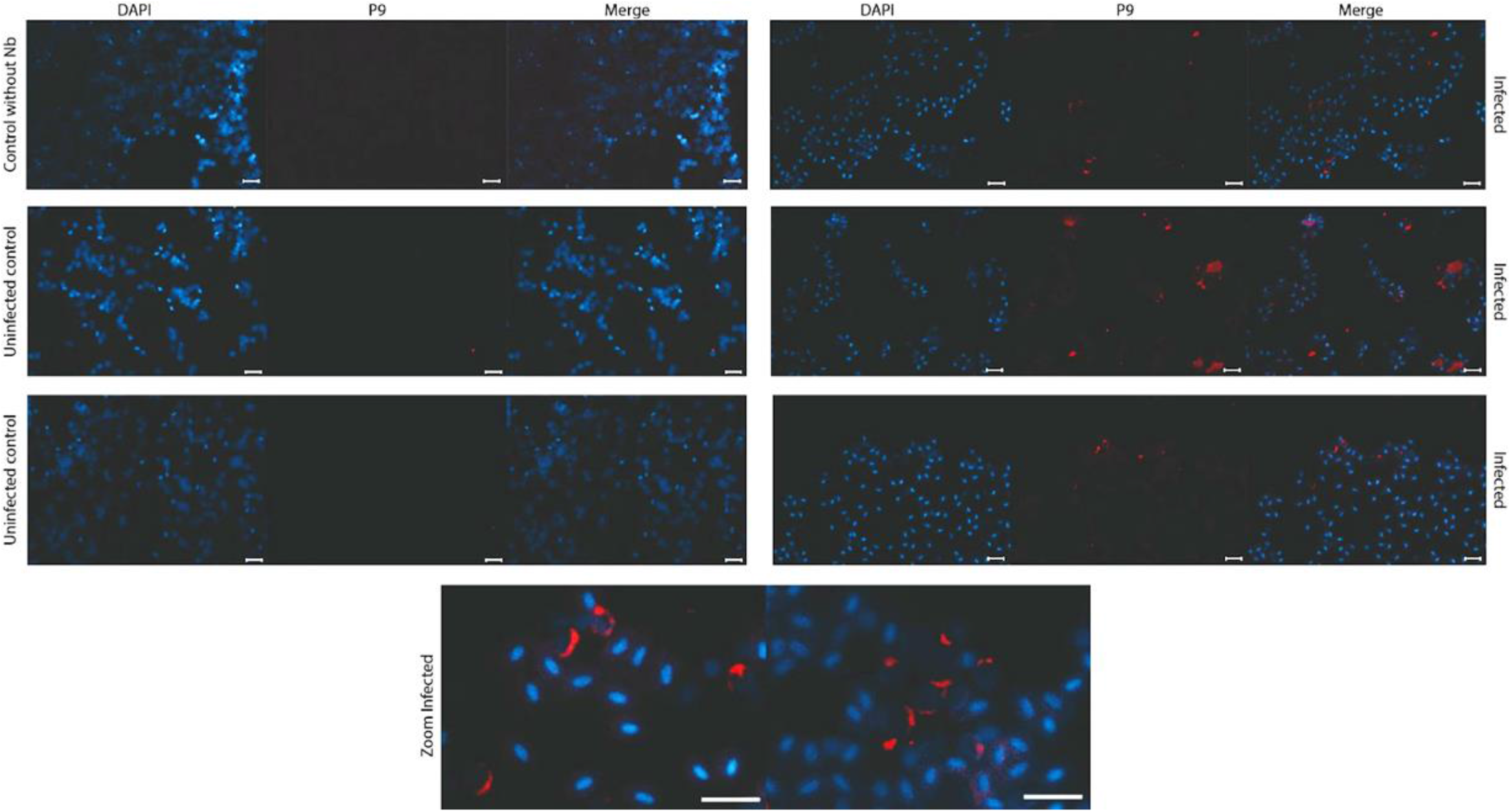
Immunofluorescence Detection of IPNv-Infected Cells in *Salmo salar*. Immunofluorescence staining of head kidney imprints from *Salmo salar* samples confirmed to be IPNv positive (by qPCR) and uninfected controls. Cells were incubated with the purified P9 nanobody as the primary antibody (1:3000 from 1mg/ml stock), followed by a mouse anti-Myc antibody and an Alexa 647-conjugated anti-mouse secondary antibody. DAPI was used to counterstain the nuclei. A distinct fluorescent signal is observed in IPNv-infected cells, indicative of specific recognition by the P9 nanobody, while uninfected control samples show no significant signal, supporting the diagnostic potential of P9 for IPNv detection in aquaculture settings. Bars=20μm

## Discussion

Nanobodies represent a promising tool for combating pathogens, highlighting the importance of enhancing and optimizing the processes by which they are generated (Valenzuela-Nieto et al., 2022). This study introduces a novel and highly efficient method for generating nanobody libraries, leveraging Type IIS restriction enzyme-based assembly to accelerate the cloning process, based on uLoop system (Pollak et al., 2020). This streamlined approach facilitated the rapid development of a bacterial display library, enabling efficient screening and selection of nanobodies. This method not only reduces the time required for library assembly but also minimizes errors associated with traditional cloning techniques. Additionally, this method enables concatenated, multi-level cloning through an ingenious design in which the ligation of the first level generates a new restriction site that can be utilized in the subsequent cloning level, allowing for rapid transfer to other vectors (Andreou and Nakayama, 2018; Casini et al., 2015; Ellis et al., 2011; Iverson et al., 2016; Lin and O’Callaghan, 2018; Romão et al., 2018). The advantages of this technology are particularly valuable in scenarios where a swift response is critical, such as during infectious disease outbreaks, shortening the time for isolation and development of specific nanobodies.

Here we have shown a successful example based on an immunization of IPNv VP2 protein. Subsequent rounds of enrichment by density gradient, using a cost-effective protocol previously developed by our group, enabled the generation of a panel of 300 nanobodies (Valenzuela Nieto et al., 2021). These nanobodies underwent secondary screening, allowing for the rapid identification of candidate clones. We report the example of clone P9, which was subsequently subcloned into a periplasmic expression vector using the *Bsa*I restriction site generated during library preparation.Thus, undoubtedly, coupling this novel and efficient library generation method with density gradient enrichment, alongside rapid high-throughput screening methods such as pooled clone ELISA or high-content microscopy, represents a significant advancement in accelerating the acquisition and identification of new nanobodies.

Our successful isolation and characterization of a nanobody (P9) with specificity for IPNv VP2 offers a contribution with the potential to pave the way for the adoption of these new technologies within the aquaculture industry. Given the significant economic impact of IPNv on the global salmon industry, this nanobody presents a promising diagnostic tool for detecting IPNv infections in fish tissues. The capacity of P9 to specifically recognize infected cells through immunofluorescence adds practical relevance, offering a potential method for viral detection in aquaculture settings. Furthermore, the development of a nanobody targeting the VP2 protein,, opens up the possibility of its therapeutic use in neutralizing viral infection, an avenue worth exploring because of its relevance for the health and sustainability of aquaculture practices.

Furthermore, the specificity and ease of production of nanobodies offer an alternative to monoclonal antibodies, which are often costly and labor-intensive to produce. Nanobodies, due to their smaller size and structural characteristics, allow for high stability and high binding affinity, making them more suitable for routine and large-scale diagnostic or therapeutic applications in the aquaculture industry (Hassanzadeh-Ghassabeh et al., 2013). The successful application of this approach to isolate a nanobody against IPNv VP2 could easily be adapted for other pathogens, broadening the impact of this methodology.

In conclusion, the nanobody library assembly technique presented here provides an innovative and efficient approach for generating high-quality nanobodies. This technology has significant potential to improve health management practices and support economic sustainability in aquaculture by addressing emerging threats, such as the appearance of new viral variants. Moreover, it is adaptable to a range of pathogens and various industrial applications.

## Acknowledgement

Felipe Serrano for illustration and Omer Navarrete for his commitment to the wellbeing of the alpacas.

---

**Supplementary Figure 1.**
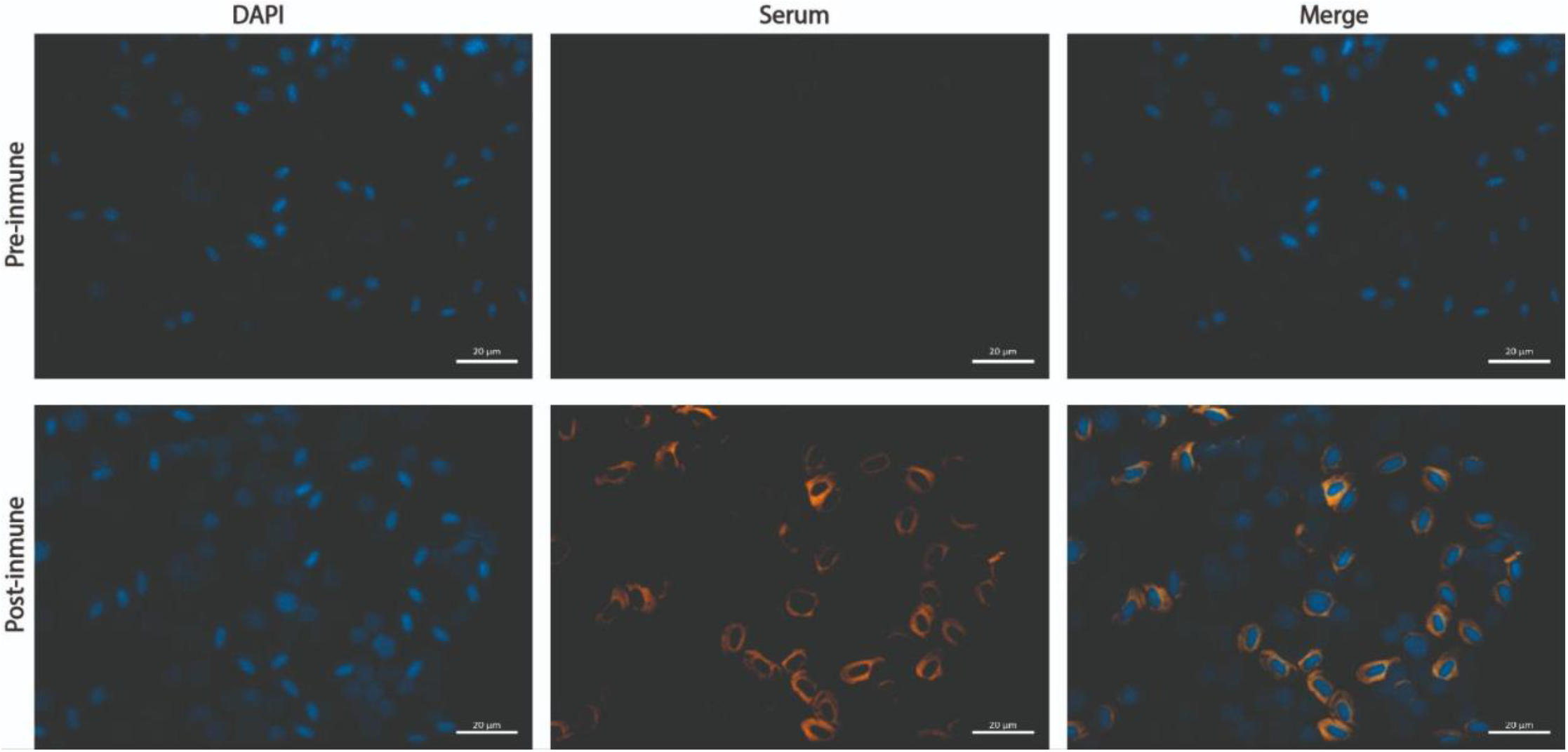
Immunofluorescence Detection of IPNv-Infected Cells in *Salmo salar* using alpaca serum. Immunofluorescence staining of head kidney imprints from *Salmo salar* samples confirmed to be IPNv positive (by qPCR) and uninfected controls. Cells were incubated with the pre and post inmunne serum as the primary antibody (1:3000), followed by a mouse anti-lama TRIT-C. DAPI was used to counterstain the nuclei. A distinct fluorescent signal is observed in IPNv-infected cells, indicative of specific polyclonal serum, while uninfected control samples show no significant signal. Bars=20μm

**Supplementary Figure 2.**
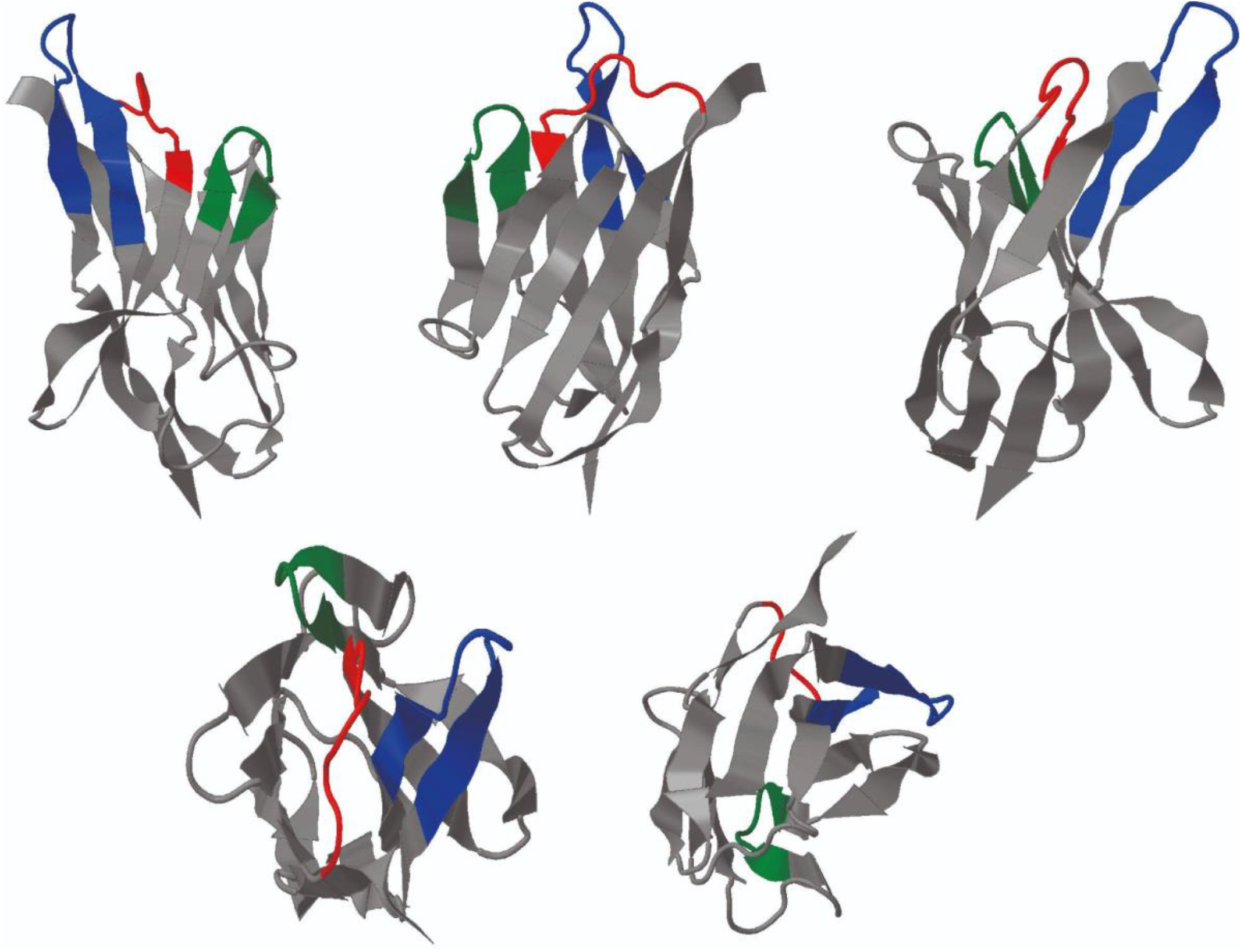
P9 nanobody structure. Alphafold structural prediction based on P9 sequence, CDRs are depicted in the same color scheme as in Fig.3

