## Supplementary figures and images for "Novel Library Assembly Technique for Developing Nanobodies Targeting IPNv VP2 Protein"

### Supplementary Figure 1

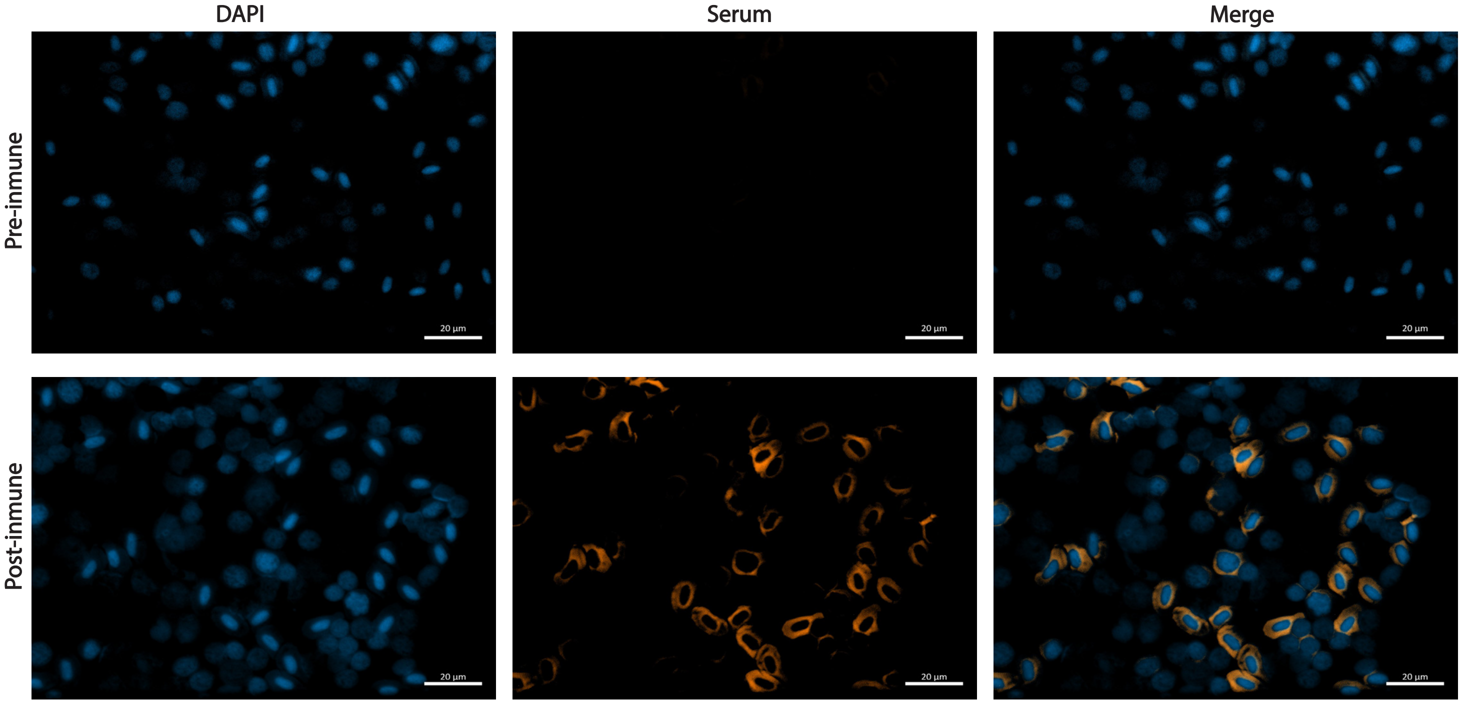

### Supplementary Figure 2

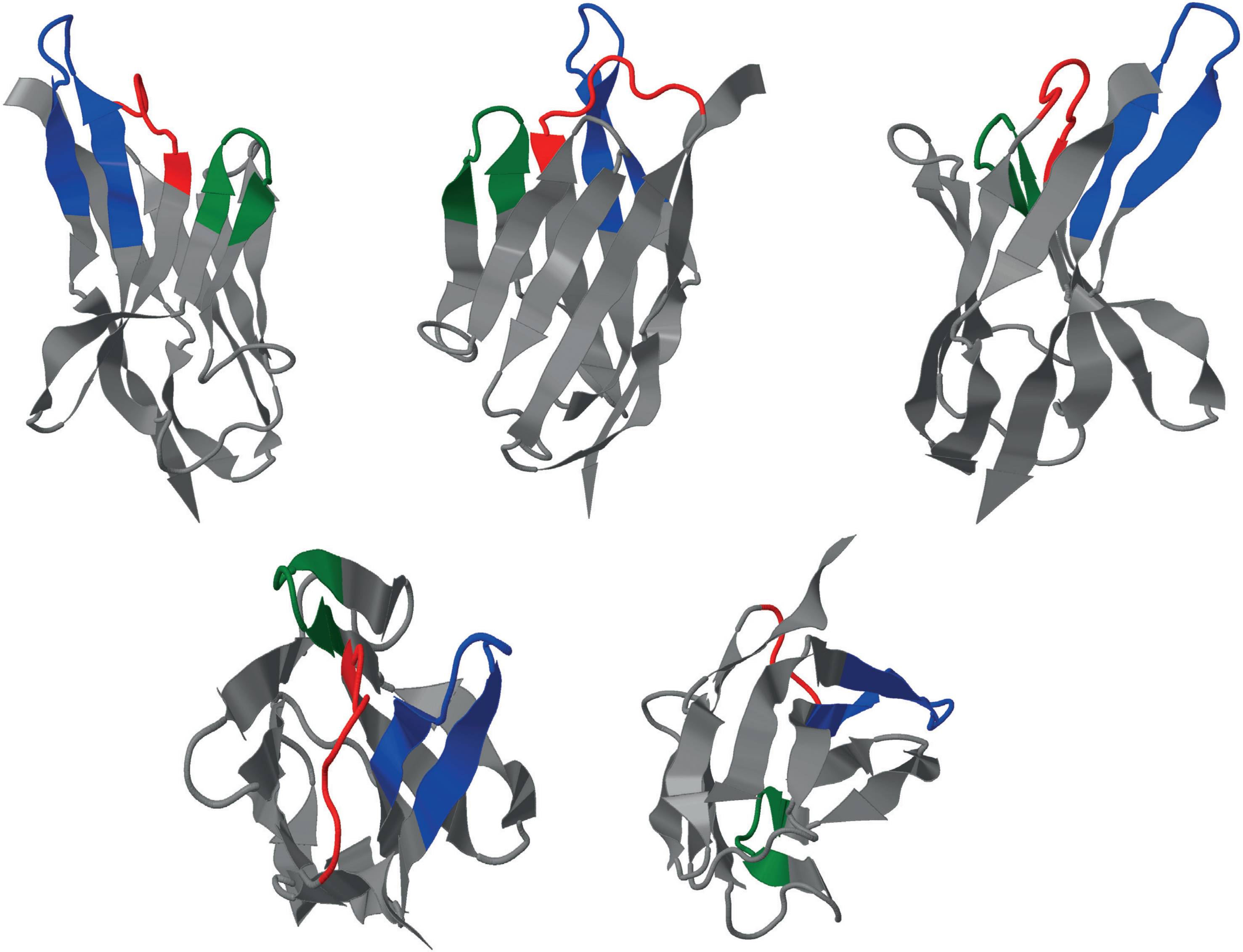
